# The best bang for the bucks: rethinking global investment on biodiversity conservation

**DOI:** 10.1101/2020.08.28.272138

**Authors:** Francisco E. Fontúrbel, Sebastián Cordero, Gabriel J. Castaño-Villa

## Abstract

Biodiversity loss is a central issue in conservation biology, being protected areas the primary approach to stop biodiversity loss. However, education has been identified as an important factor in this regard. Based on a database of threatened species and socio-economic features for 138 countries, we tested whether more protected areas or more education investment are associated with a lower proportion of threatened species (for different groups of vertebrates and plants). We found that education investment was negatively associated with the proportion of threatened species in 2007 and 2017, as well as with their change rates. Conversely, the percentage of protected land was significant for reptiles, but show weak relationships with other groups. Our results suggest that only increasing protected areas will not stop or reduce biodiversity loss, as the context and people’s attitudes towards wildlife also play major roles here. Therefore, investing in education, in addition to protected areas, would have a positive effect missing to achieve effective species conservation actions worldwide.

## 1. Introduction

Biodiversity loss is the most central issue in conservation biology. As a result of human actions, hundreds of species went extinct during the last centuries (Humphreys *et al*., 2019), and hundreds more are likely to be extinct in the short term. In response to this global problem, the United Nations created the Convention on Biological Diversity (CBD) in 1992, seeking a multi-lateral agreement to achieve sustainable development and stop biodiversity loss (Balmford *et al*., 2005). Since then, we acknowledged that the solution to stop biodiversity loss not only relies on the academy and the scientific knowledge but also on social and economic factors (Hooper *et al*., 2012). Therefore, a multi-dimensional approach is needed to deal with biodiversity loss crisis at both local and global scales (Newbold *et al*., 2015). In this regard, the United Nations 2030 Agenda for Sustainable Development points education and protected areas as key instruments to reduce biodiversity loss and improve people’s life quality.

Most countries in the world have improved and expanded their protected area systems in order to slow biodiversity loss and create conservation spaces in which the extant biodiversity could be spared (Butchart *et al*., 2010; Cardinale *et al*., 2012). However, the effectiveness of protected areas is largely debated (Rodrigues *et al*., 2004; Carranza *et al*., 2014; Watson *et al*., 2014). The increase of protected area extent across countries seems not to be enough to reduce biodiversity loss. Those spare lands intended as biodiversity preservation spaces are not independent of their context (i.e., many protected areas are surrounded by a heterogeneous matrix of disturbed lands), sometimes are not large enough, and in many cases are not representative enough (Armesto *et al*., 1998; Durán *et al*., 2013). As protected surface increases, the human population, urban areas, and the percentage of the human population living in cities also increase, posing a major threat to the biodiversity outside protected area borders (Mcdonald *et al*., 2008; Sanderson *et al*., 2018).

An important issue within the socio-economic dimension of biodiversity conservation is education. In this regard, a lack of conservation opportunities is associated with poor education on countries within biodiversity hotspots (Acosta, 2000). In this regard, education (and particularly education quality) is acknowledged as a key (but not quite evident) factor determining positive attitudes towards biodiversity (Caro *et al*., 1994). Therefore, education for conservation is amongst the top objectives of conservation biology, as it could be the necessary force to achieve long-term society changes, making attitudes to nature more positive (Strauss, 1995).

Here, we present a global assessment of the relative importance of protected areas and education investment on the proportion of threatened vertebrate and plant species. Based on socio-economic and biological information from 138 countries from the seven continents (Africa, Asia, Australia, Europe, Central American, North America, South America), we assessed the relative importance of protected areas and education (which are two major investment sectors for biodiversity conservation) on explaining the proportion of threatened species in 2007 and 2017, as well as the change in threat levels in these ten years. We tested the following hypotheses: (1) if protected areas are sufficient by themselves, larger protected area surfaces and larger proportions of land spared as protected lands should be associated with lower proportions of threatened species; (2) More investment in education should generate positive attitudes towards biodiversity, and therefore we expect lower proportions of threatened species in those countries investing more money on education.

## 2. Material and methods

### 2.1 Data sources

We built two datasets. The first dataset contains information regarding the proportion of threatened species of terrestrial vertebrates (i.e., amphibians, reptiles, birds, mammals, and all combined) and plants for two time periods: 2007 and 2017. We obtained the number of species per group present at each country from the following databases: eBird 2017 edition (https://ebird.org) and Avibase 2017 edition (https://avibase.bsc-eoc.org) for birds, The Reptile Database 2017 edition (http://reptile-database.reptarium.cz/) for reptiles, Amphibians Species of the World (Frost, 2017) for amphibians, Mammal Species of the World (Wilson and Reeder, 2005), and the Guide to Standard Floras of the World (Frodin, 2010) for vascular plants, supplemented with regional databases of countries not included in the list. Additionally, we obtained the number of threatened species per group from the IUCN redlist database (http://www.iucnredlist.org).

The second dataset contains socio-economic information of each country (Table S1), composed by the following variables: geography (mainland or island), country size (in km^2^), population size, population density (hab./km^2^), urbanization (% of the country area corresponding to urban areas), Gross Domestic Product (GDP hereafter, expressed in M USD), % of the GDP invested in education, USD invested in education (resulting from the multiplication of the last two variables), protected areas extent (km^2^), and the % of the country designed as a protected area (resulting from dividing protected area surface by country size * 100%). We selected those variables as they represent most of the aspects of human actions over native flora and fauna (i.e., we expect urbanization and population density to have a negative impact on biodiversity, whereas we expect protected areas to have a positive impact in turn), and also give context (i.e., threatened species may depend if a given country is an island or not, country size, and people’s education level and income may also affect the ways to relate with wildlife; Kier *et al*., 2009). We obtained country information (corresponding to data from 2006) from the CIA Factbook website (https://www.cia.gov/library/publications/the-world-factbook/). Although there is updated information, we used the 2006 dataset to account for the time lag in education effects (i.e., the effects of investing in education are not immediate). We gathered information for 138 countries across the world (we did not include countries with incomplete information).

### 2.2 Data analysis

First, we conducted a correlation analysis among the socio-economic variables (Fig. S1) in order to detect highly correlated and potentially multicollinear variables, aiming to avoid variable redundancy in the models. Then, we transformed the proportion of threatened species for each group and year using the following formula:

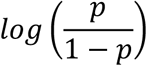

We transformed the percentage of protected area per country using log (*p* + 1). We used these transformations to improve data following a Gaussian distribution.

We fitted Mixed-effects Generalized Linear Models (GLMM; Zuur *et al*., 2009), using the proportion of threatened species as the response variable. We included the following socio-economic variables as predictors (after filtering for redundant variables): % of GDP invested in education, protected surface, and % of protected area. We included urbanization and population density as covariates (in order to account for major differences among countries). Geography (mainland or island) was included as a random factor to account for the geographic variability, as islands tend to have higher threat levels than mainland countries due to their biogeographic context (Kier *et al*., 2009). GLMM parameters and their significance were estimated using restricted maximum likelihood t-tests with a Kenward-Roger approximation to degrees of freedom (Halekoh and Højsgaard, 2014). Then, we used multi-model inference (Burnham and Anderson, 2004) to assess the relative importance of each predictor variable within the GLMM models fitted. For this, we obtained the delta ≤ 2 model subset based on the BIC rank. The resulting barplot represents the percentage of models of the ΔBIC ≤ 2 subset in which each predictor variable occurs, ordered from the most important to the least (Burnham and Anderson, 2004). We used this procedure to examine 2007 and 2017 datasets, and also to examine the change between 2017 and 2007 (i.e., creating new response variables from the difference between 2017 and 2007 datasets).

Then, to examine the significance of the change in threatened species proportion between 2007 and 2017, we used paired *t*-tests for each group. Finally, we contrasted our 2006 protected area percentages per country with the most recent data from the World Database on Protected Areas (WDPA, www.protectedplanet.net, updated June 2019) to estimate the increase in protected area coverage at each country between 2006 and 2019, aiming to relate such increase with the change in threatened species proportions. All analyses were conducted in R 3.5.1 (R Development Core Team, 2018), using the packages *mgcv* (Wood and Scheipl, 2014), *lme4* (Bates *et al*., 2013), *lmerTest* (Kusnetzova *et al*., 2015), *pbkrtest* (Halekoh and Højsgaard, 2014), and *MuMIn* (Bartoń, 2018).

### 2.3 Data accessibility

Species and full socio-economic databases, the R code used, and its detailed outputs are available from the *figshare* digital repository (doi: 10.6084/m9.figshare.9171992).

## 3. Results

Examining the effects of the socio-economic variables on the proportion of threatened species, we found for 2007 that education investment was significant for vertebrates combined (and particularly for reptiles and amphibians), while for birds and mammals it was not significant, showing that countries that invest more money (i.e., a more significant percentage of the GDP) have lower proportions of threatened species. Conversely, the percentage of protected land was significant only for reptiles, and the protected surface was significant for birds. However, this relationship is positive, suggesting that countries with more protected areas also have more threatened bird species (Table 1a). Examining the 2017 dataset, education presented significant effects on vertebrates (and particularly on mammals and reptiles) and plants, showing that more investment in education is associated with less threatened species. The percentage of protected land was significant for mammals, reptiles, and all vertebrates, whereas the protected surface was significant for birds but with a positive relationship as in 2007 (Table 1b). In all cases, the proportion of threatened species reduces with education investment and the percentage of protected land, but it increases with the protected surface, particularly on birds. Examining the relative importance of the socio-economic variables, we found that education investment is the most important predictor in all cases, with support values ranging between 50 and 100%. The next variable in importance is the percentage of protected land, which was important for birds, reptiles, and amphibians. Protected surface and population density were the least important variables in all cases (Fig. S2).

**Table 1.**
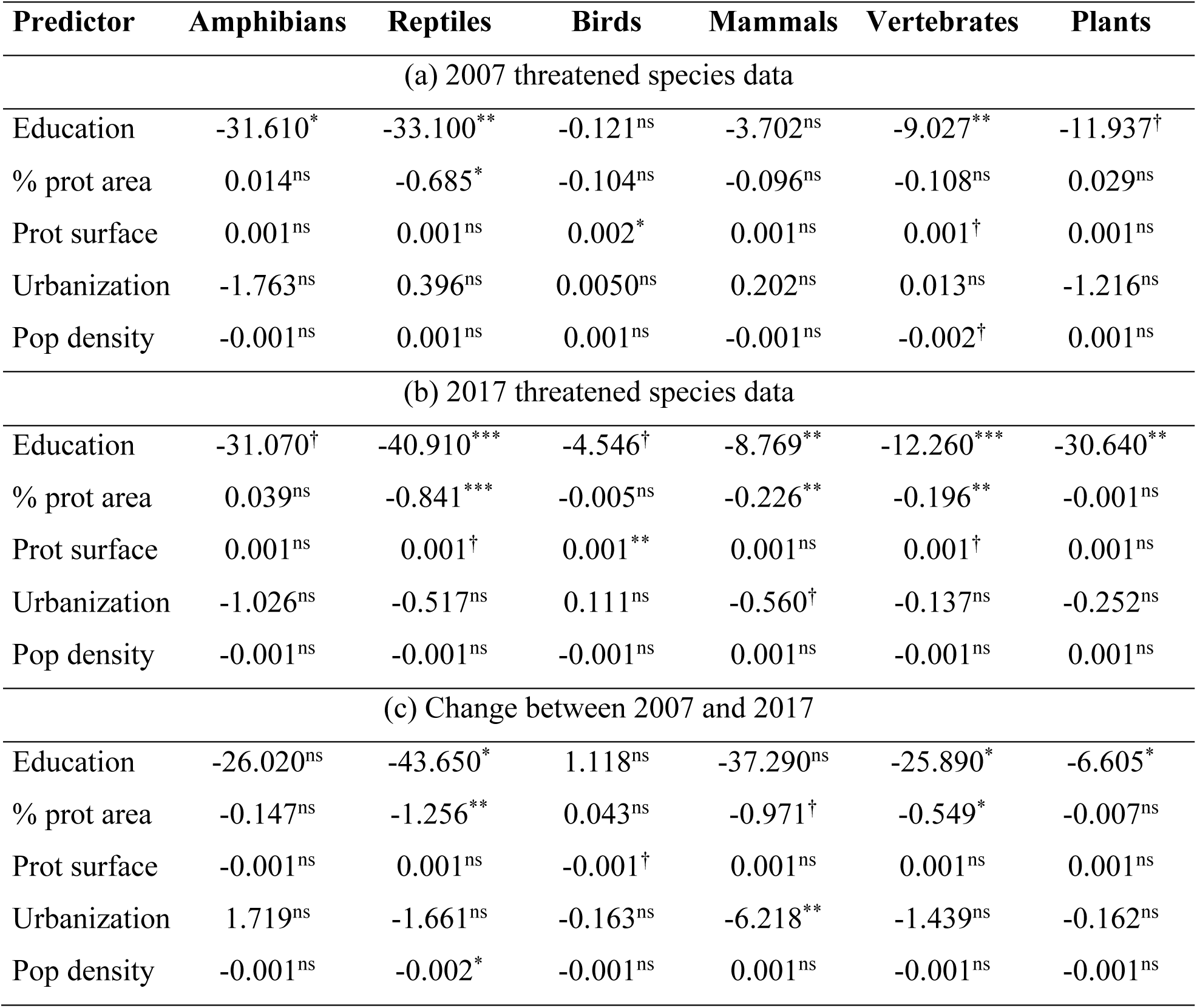
Summary of the effects of socio-economic predictors on the proportion of threatened species in (a) 2007, (b) 2017, and (c) the change between 2007 and 2017. For each case *t*-values and their significance are presented (ns = *P* > 0.10; † = 0.05 < P < 0.10, * = P < 0.05, ** P < 0.01, *** = P < 0.001).

Comparing the proportion of threatened species between 2007 and 2017, we found that the proportion of threatened mammals was not significantly different. However, the proportion of threatened birds, reptiles, amphibians, total vertebrates, and plants have significantly increased in these ten years (Table 2). Mammals, reptiles, and amphibians had the largest variation ranges, with contrasting changes across countries. Regarding the effects of socio-economic variables on the change rates, education was significant for vertebrates (mainly reptiles) and plants. In contrast, the proportion of protected areas was significant for vertebrates and particularly reptiles (Table 1c). Examining the relative importance of these variables in the change of threatened species proportion, education had the largest support across models, followed by the percentage of protected land, which was important for reptiles and amphibians (Fig. S3).

**Table 2.** Comparison of the proportion of threatened species of mammals, birds, reptiles, amphibians, all vertebrates, and plants between 2007 and 2017, change rate (Δ = 2017 – 2007), and variation range (minimum and maximum change values). We assessed statistical differences using a paired *t*-test (two-tailed).

| Comparison | % 2007 | % 2017 | Change | Range | *t* | df | *P* |
| --- | --- | --- | --- | --- | --- | --- | --- |
| Amphibians | 9.76% | 10.58% | 0.83% | -11 to 27 % | -2.417 | 137 | 0.017 |
| Reptiles | 4.19% | 7.53% | 3.32% | -8 to 33 % | -6.503 | 137 | <0.001 |
| Birds | 2.61% | 3.67% | 1.00% | -1 to 4 % | -18.545 | 137 | <0.001 |
| Mammals | 10.74% | 11.49% | 0.72% | -7 to 44 % | -1.500 | 137 | 0.136 |
| Vertebrates | 6.85% | 8.33% | 1.49% | -3 to 25 % | -5.731 | 137 | <0.001 |
| Plants | 0.79% | 1.17% | 0.28% | 0 to 4 % | -6.339 | 136 | <0.001 |

Examining changes in threatened species proportions between 2007 and 2017, we found that all groups assessed here have experienced an increase in the proportion of threatened species between 2007 and 2017 (Table S2). However, in some cases, the differences among countries are considerable (for example, change rates for mammals range from −7% to 44%). Contrasting those rate changes to our socio-economic variables, we found that countries investing more resources in education (expressed as the percentage of the GDP invested in education in 2006, with ten years of time lag), have lower or even negative change rates over these ten years. This pattern was consistent across groups except for birds (cf. Fig. 1). Then, examining its relationship with the percentage of protected land, we found weak relationships in all cases (Fig. S4). Finally, examining the relationship between the change of threatened species between 2007-2017 and the increase in protected area surface between 2006 and 2019, we observe contrasting results among groups (cf. Fig. 2). However, none of those relationships was statistically significant (Table S3).

**Figure 1.**
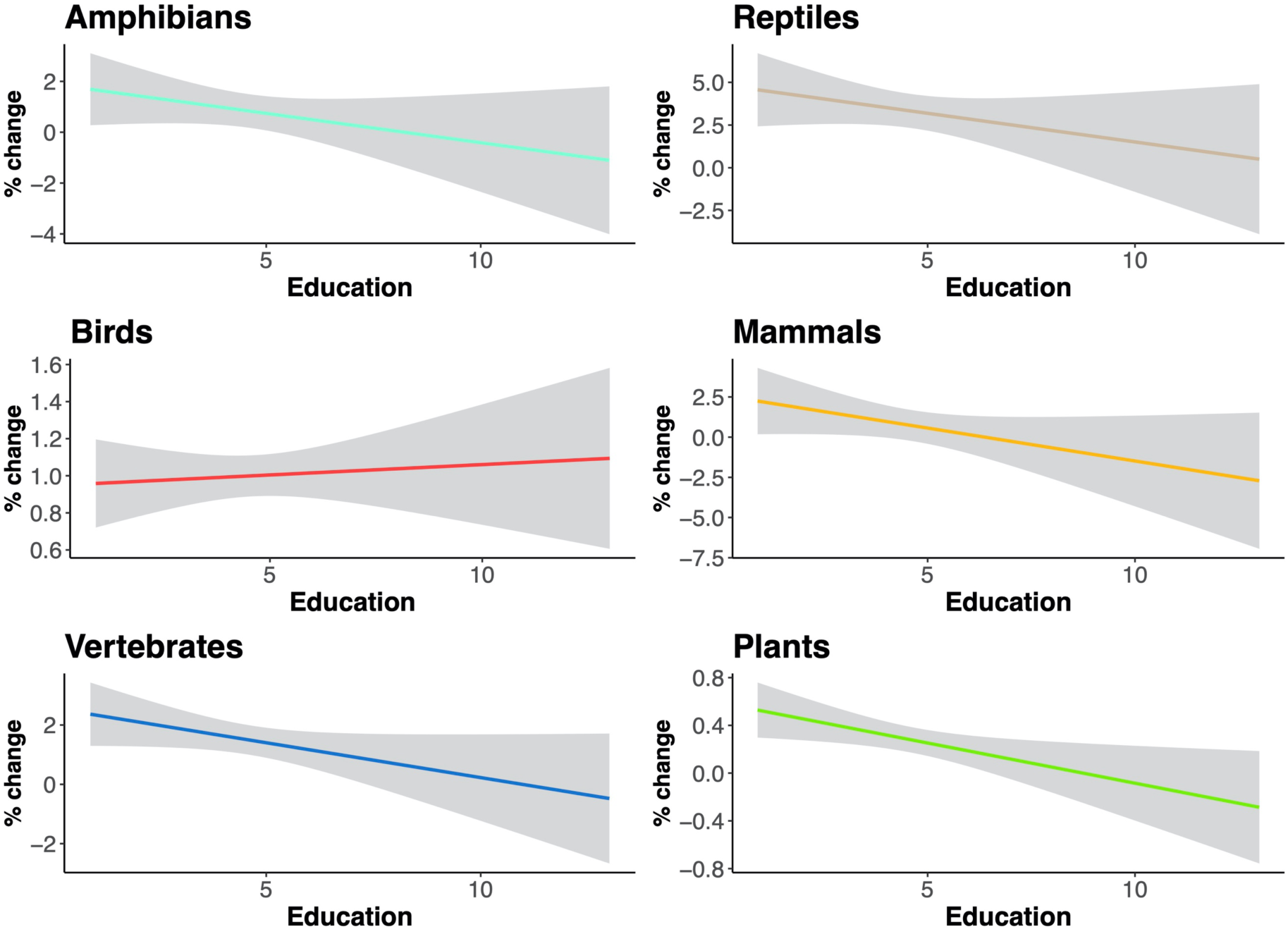
Relationships between education investment (expressed as the percentage of GDP invested in education) and the change in the proportion of threatened species between 2017 and 2007. Gray areas represent the 95% confidence interval.

**Figure 2.**
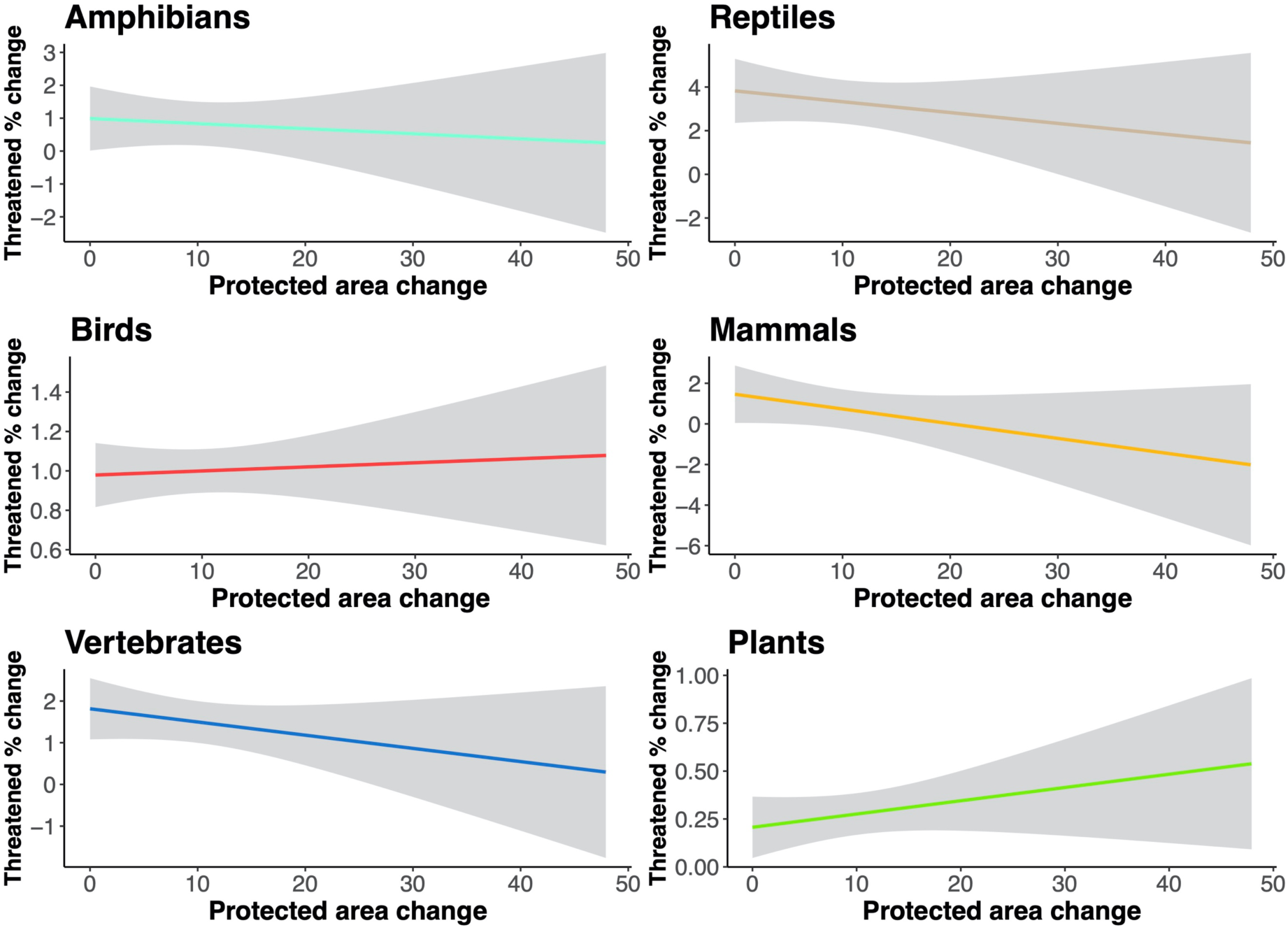
Relationships between protected area increase between 2006 and 2019 (based on the percentage of each country designated as protected area) and the change on the proportion of threatened species between 2017 and 2007. Gray areas represent the 95% confidence interval.

## 4. Discussion

Our results point out that investment in education can be as crucial as sparing land as protected areas for wildlife conservation. Those countries investing more money in education (see Figs. S5 and S6) consistently had lower proportions of threatened vertebrates and plants. The most substantial result of our assessment is the significant effect of education investment on the proportion of threatened vertebrate and plant species, as well as on their change in ten years. Thus, education plays a central role in biodiversity conservation as it generates positive attitudes towards wildlife (Caro *et al*., 1994), empowers people, and with the appropriate information, encourages them to take action. Such attitude changes are essential to change the conservation paradigm and achieve changes in the long-term (Strauss, 1995; Pyke *et al*., 1999). Countries that invest larger proportions of their GDP (irrespective of the amount of such GDP) in education may have more opportunities to achieve attitude changes in the population. This outcome is evident from our dataset as the money invested in education in 2006 showed a favorable effect on threatened species in 2017. Educative processes involve changes in the mid- and long-term, as the resources invested now to improve education quality will show their effects a decade later or more as a result of a progressive attitude change (particularly on children and young people), as the Sustainable Development 2030 Agenda proposes.

Despite its importance to conservation practice, protected areas alone have a limited effect on relieving biodiversity loss. Designating protected areas not only a scientific process as it also involves political decisions. Therefore, the weak effect of increasing protected areas on reducing the proportion of threatened species could be related to the common practice of protecting marginal lands instead of high priority areas (Armesto *et al*., 1998; Duarte *et al*., 2014). For example, Chile has protected ∼22% of its territory, but 89% of those protected areas are located in latitudes > 43°S, resulting in an uneven distribution of the area under protection regime (Armesto *et al*., 1998), leaving many conservation-concern and highly threatened taxa unprotected (e.g., Duarte *et al*., 2014). In this example, most of the threatened and endemic species, along with specialized ecological interactions occur in the Mediterranean type-ecosystem of central Chile (Medel *et al*., 2018), which has the lowest protected area coverage, and also is highly-threatened by the expansion of urban areas, vineyards, and avocado plantations. If we examine this pattern at the global scale, we observe a moderate correspondence of protected areas with biodiversity hotspots (Fig. 3), which is even lower when we examine threatened species instead (Fig. 4). For example, the Brazilian Atlantic rainforest, the Indian peninsula, and the south-east Asia archipelago have the largest proportions of threatened birds, but protected areas are negligible at those locations (Peh *et al*., 2006; Barlow *et al*., 2007; Bohada-Murillo *et al*., 2020). A similar pattern is observed for mammals of south-east Asia and amphibians of eastern China. From the keystone work of Margules and Pressey (2000), systematic conservation planning has oriented decision-making regarding protected areas. In this regard, protected areas alone are not enough to dampen biodiversity loss, as their success depends on landscape and socio-economic contexts, which should also be taken into account.

**Figure 3.**
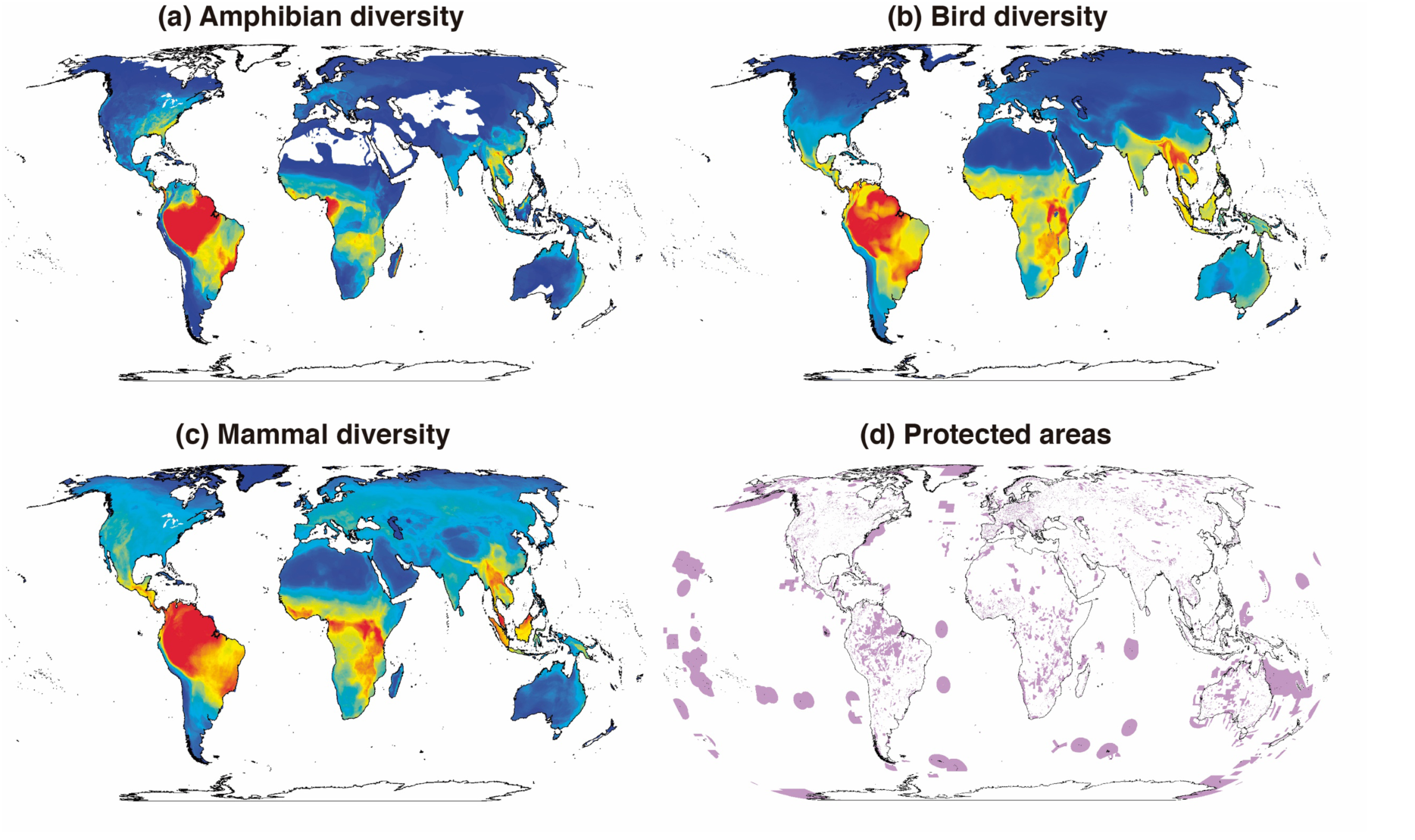
Global species richness of (a) amphibians, (b) birds, and (c) mammals. World protected areas are shown in panel (d). Diversity data was obtained from Jenkins *et al*. (2013) and Pimm *et al*. (2014). Protected area data was obtained from the World Database on Protected Areas (WDPA, www.protectedplanet.net, updated June 2019).

**Figure 4.**
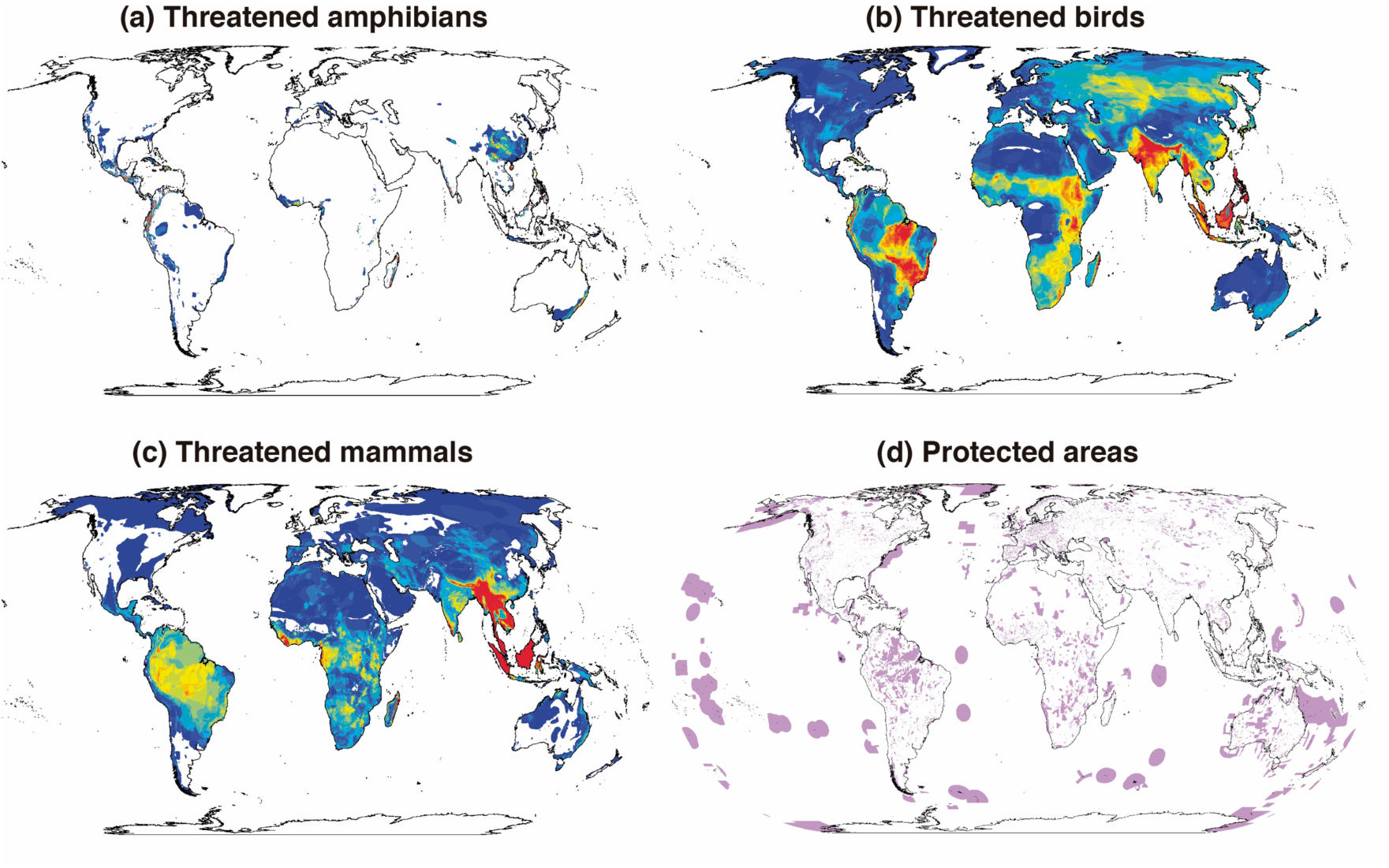
Globally threatened species of (a) amphibians, (b) birds, and (c) mammals. World protected areas are shown in panel (d). Diversity data was obtained from Jenkins *et al*. (2013) and Pimm *et al*. (2014). Protected area data was obtained from the World Database on Protected Areas (WDPA, www.protectedplanet.net, updated June 2019).

Also, the exponential increase of the human population and their rapid concentration on urban areas are strong drivers of biodiversity loss. We included population density and urbanization as covariates in our models because they have a negative correlation with biodiversity across taxonomic groups but is particularly critical for reptiles and amphibians (Hamer and McDonnell, 2010; French *et al*., 2018). On the other hand, plant diversity is also negatively affected by urbanization and population densification (Goddard *et al*., 2010), exerting adverse impacts on the structure and composition of plant communities, and leading to the extinction of native plants and the spread of invasive species (Hahs *et al*., 2009). These impacts can trigger a cascade effect on biodiversity loss since the loss of plant species usually results in the loss of mammal species associated with the vegetation structural features (McKinney, 2008). Large mammals, in particular, are expected to be severely affected by urbanization as they require large habitat extensions, which become a limiting resource as a result of urbanization processes (McKinney, 2008), and are the primary target of protected area-based conservation. In any case, native populations tend to reduce under extreme urbanization conditions, irrespective of the taxonomic group (McKinney, 2008).

## 5. Conclusion

Stopping biodiversity loss is a complex and challenging task that requires a multi-dimensional approach. Our results point to education as the primary change force that can trigger a long-term attitudinal change towards wildlife and its conservation. Without people’s involvement, any other effort may have limited success. Even though protected areas comprise about 12% of the terrestrial surface, global biodiversity continues progressively declining (Butchart *et al*., 2010). Those protected areas are suffering biodiversity erosion processes as a result of habitat disturbance, hunting, defaunation, and overexploitation (Dirzo *et al*., 2014), which are directly related to people’s attitudes and their access to information. Despite protected areas being effective, avoiding deforestation within its boundaries, what happens outside them largely depends on people’s attitude, and those attitudes are ultimately influenced by education. Protected areas are essential for biodiversity conservation, with no doubt (Leverington *et al*., 2010), but we cannot rely only on them. We urge a paradigm change, not only prioritizing high diversity areas, but also shifting to an integrative framework beyond protected areas including areas with endangered / endemic species, along with adequately representing the diversity of the ecosystems, including the people, and controlling the rapid expansion of urban settlements over natural areas with high-level policies.

## Acknowledgements

Comments of J.A. Simonetti on an earlier version of this manuscript helped us to clarify our initial ideas and focus on the analysis. We received no funding for this study.

## Supplementary Data

List of the countries included in our assessment (Table S1), correlation matrix among socio-economic variables (Fig. S1), the relative importance of the variables included in the GLMM models (Figs. S2 and S3), relationships between protected land and change on threatened species (Fig. S4), changes in threatened species proportions between 2007 and 2017 (Table S2), and detailed GLMM results of the relationship between the percentage of threatened species and the increase on protected areas (Table S3) are available online as Supplementary Material.

