## Supplementary Data for "The best bang for the bucks: rethinking global investment on biodiversity conservation"

**Table S1.** List of the 138 countries from which we obtained socio-economic information, ordered by continent and geographical context (mainland or island).

| Continent | Mainland countries | Island countries |
|---|---|---|
| Africa | Algeria, Angola, Benin, Botswana, Burkina Faso, Burundi, Cameroon, Central African Republic, Chad, Congo, Cote d'Ivoire, Djibouti, Egypt, Eritrea, Ethiopia, Gabon, Gambia, Ghana, Guinea, Kenya, Lesotho, Libya, Malawi, Mali, Mauritania, Mozambique, Namibia, Niger, Nigeria, Rwanda, Senegal, Sierra Leone, South Africa, Sudan, Swaziland, Tanzania, Togo, Tunisia, Uganda, Zambia, Zimbabwe | Madagascar, Mauritius |
| Asia | Armenia, Azerbaijan, Bangladesh, Bhutan, Cambodia, China, Georgia, India, Iran, Israel, Jordan, Kazakhstan, Korea, Kuwait, Lebanon, Malaysia, Mongolia, Nepal, Oman, Pakistan, Russia, Saudi Arabia, Thailand, Turkey, Turkmenistan, Uzbekistan, Viet Nam, Yemen | Indonesia, Japan, Philippines, Singapore |

| | | |
|---|---|---|
| Europe | Albania, Austria, Belarus, Belgium, Bulgaria, Croatia, Czech Republic, Denmark, Estonia, Finland, France, Germany, Greece, Hungary, Italy, Latvia, Lithuania, Macedonia, Moldova, Netherlands, Norway, Poland, Portugal, Romania, Slovakia, Slovenia, Spain, Sweden, Switzerland, Ukraine | Iceland, Ireland, United Kingdom |
| Oceania | (none) | Fiji, New Zealand, Australia |
| South America | Argentina, Bolivia, Brazil, Chile, Colombia, Ecuador, Guyana, Paraguay, Peru, Uruguay, Venezuela | (none) |
| Central America | Belize, Costa Rica, El Salvador, Guatemala, Honduras, Nicaragua, Panama | Barbados, Cuba, Dominican Republic, Haiti, Jamaica, Trinidad and Tobago |
| North America | Canada, Mexico, United States | (none) |

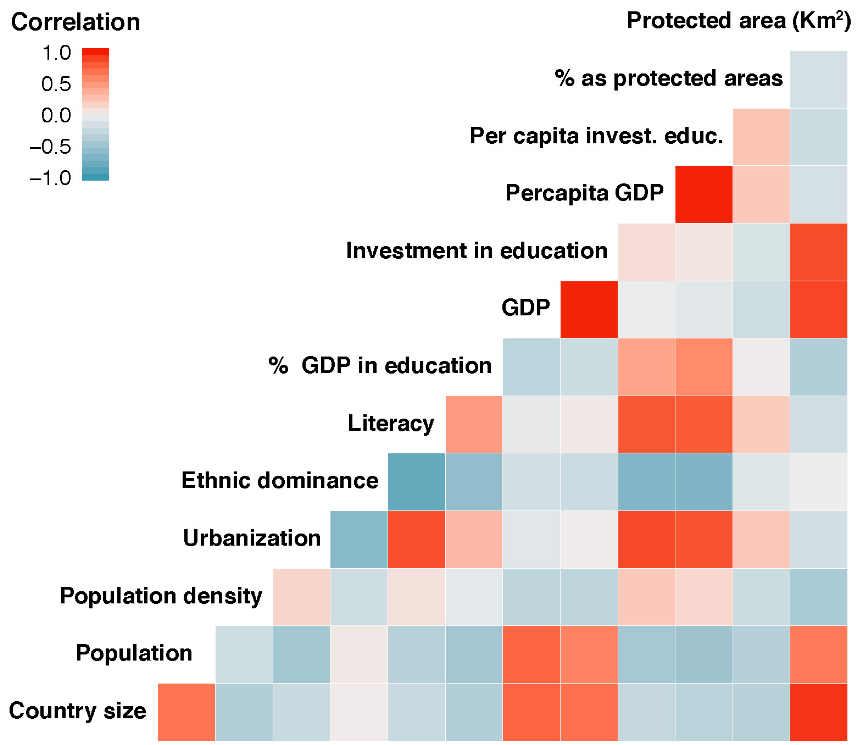


**Figure S1.** Correlation among socio-economic variables included in the original database. GDP stands for Gross Domestic Product. Source: CIA Factbook (data from 2006).

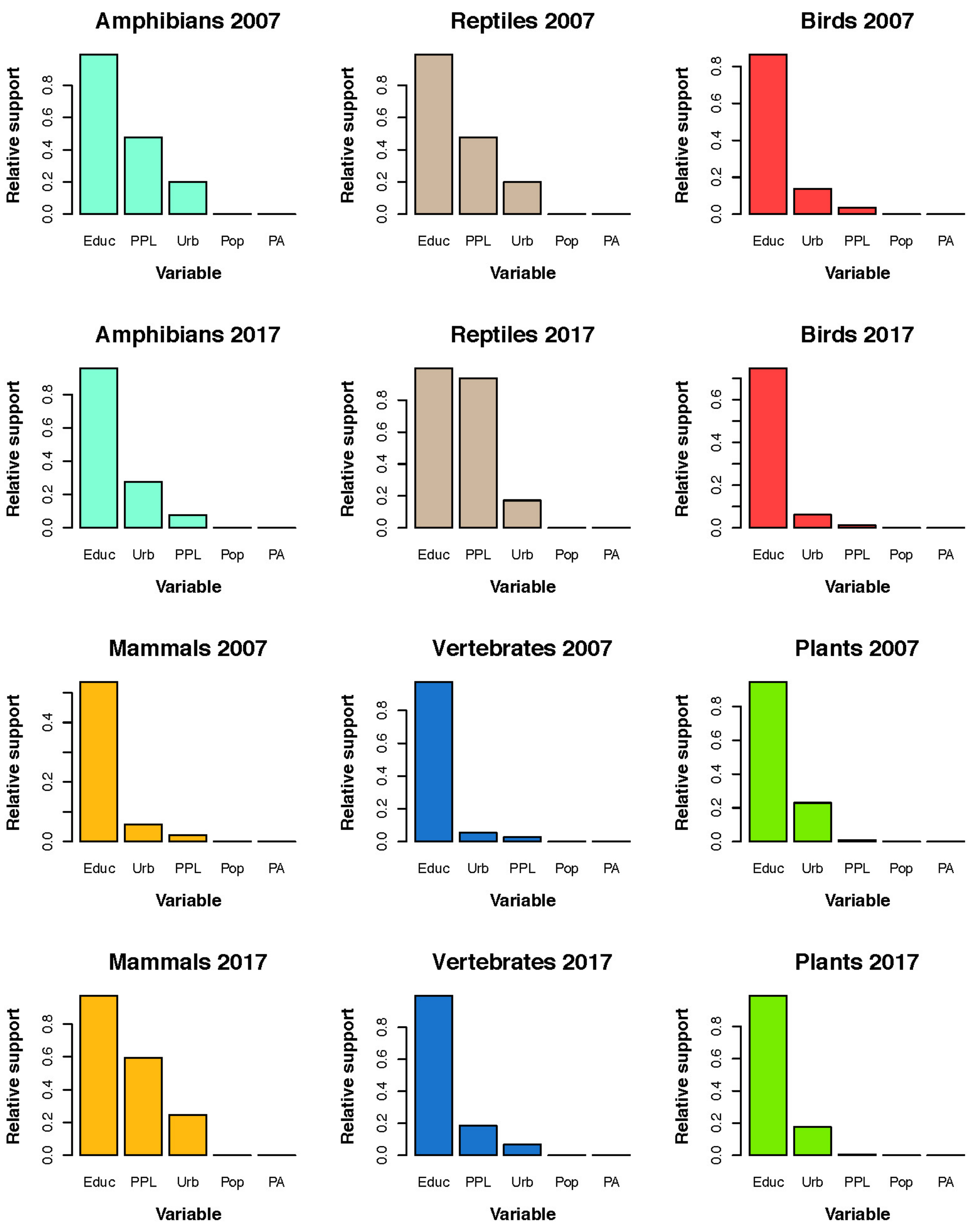


**Figure S2.** Relative importance of the variables included in the GLMMs fitted for threatened species proportions in 2007 and 2017. Abbreviations: Educ = education investment, PPL = percentage of protected land, PA = protected area surface, Urb = urbanization, Pop = population density.

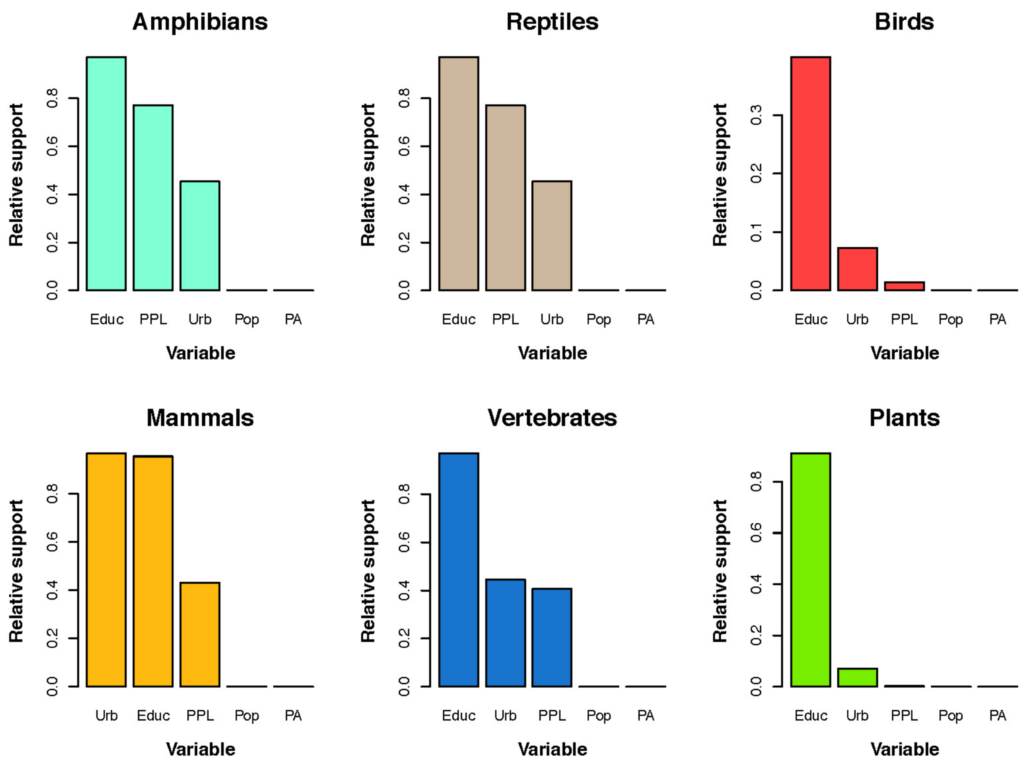


**Figure S3.** Relative importance of the variables included in the GLMMs fitted for threatened species change between 2007 and 2017. Abbreviations: Educ = education investment, PPL = percentage of protected land, PA = protected area surface, Urb = urbanization, Pop = population density.

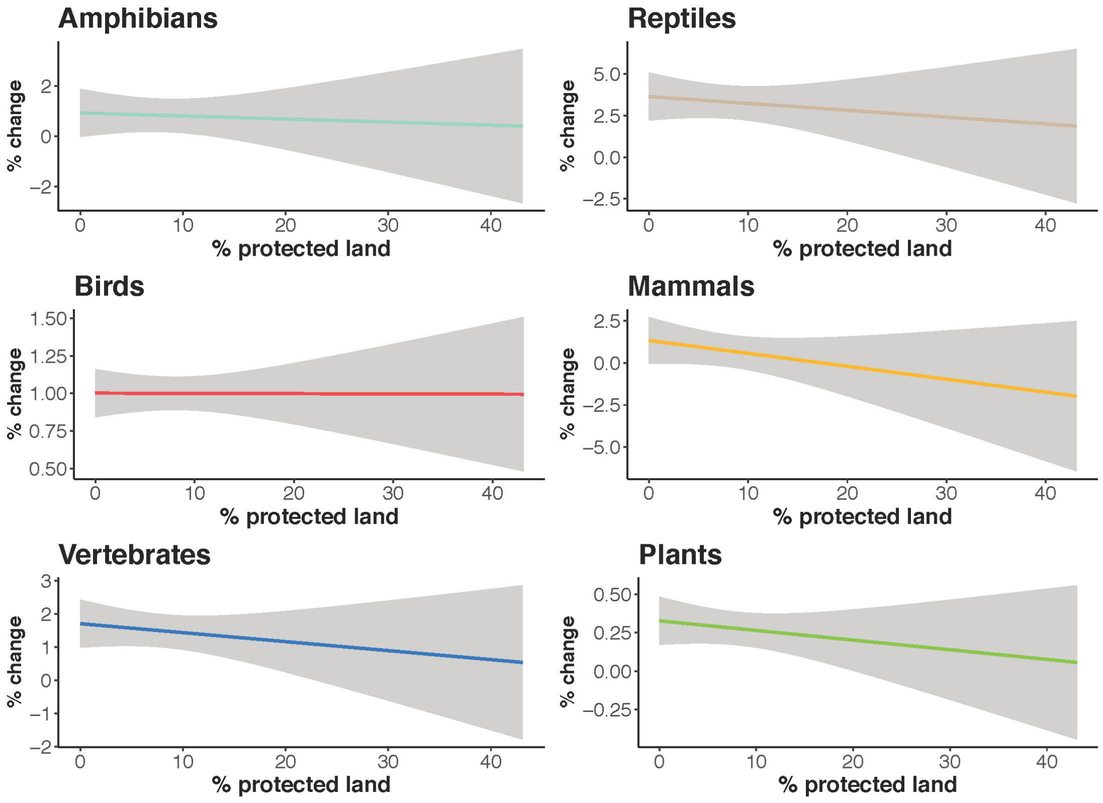


**Figure S4.** Relationships between the percentage of country surface designated as protected area (data from 2006) and the change on the proportion of threatened species between 2017 and 2007.

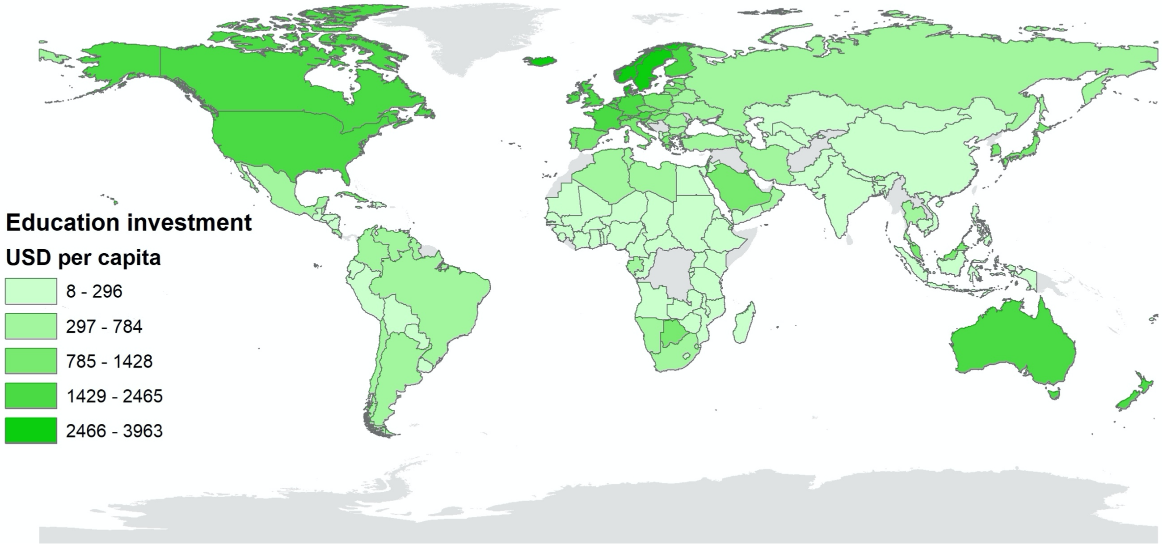


**Figure S5.** Education investment map, expressed in US dollars per capita per year, for the 138 countries included in our database. Gray areas represent no data available.

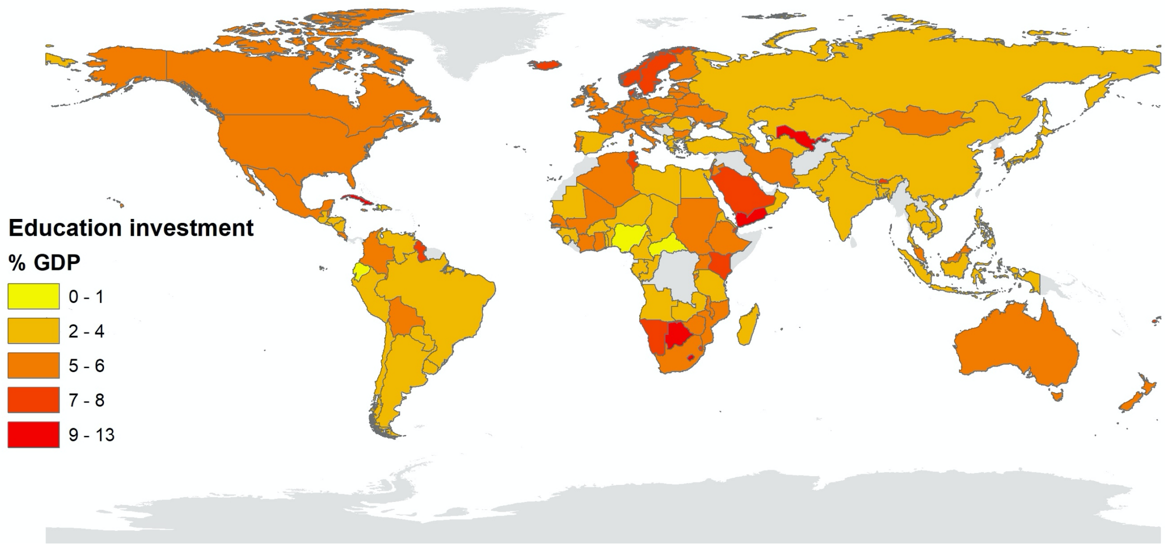


**Figure S6.** Education investment map, expressed in % of the GDP, for the 138 countries included in our database. Gray areas represent no data available.

**Table S2.** Proportion of threatened species by group for 2007 and 2017, their change in a 10-year period, and the delta range among countries.

| Group | Prop. 2007 | Prop. 2017 | Delta | Range |
|---|---|---|---|---|
| Amphibians | 9.76% | 10.58% | 0.83% | -11 to 27 % |
| Reptiles | 4.19% | 7.53% | 3.32% | -8 to 33 % |
| Birds | 2.61% | 3.67% | 1.00% | -1 to 4 % |
| Mammals | 10.74% | 11.49% | 0.72% | -7 to 44 % |
| Vertebrates | 6.85% | 8.33% | 1.49% | -3 to 25 % |
| Plants | 0.79% | 1.17% | 0.28% | 0 to 4 % |

**Table S3.** Detailed results of the GLMM models fitted to assess the relationship between the change in threatened species per group over a 10-year period (2007-2017) and the change in surface designated as protected area between 2006 and 2019.

| Group | Estimate | SE | t value | P value |
|---|---|---|---|---|
| Amphibians | -0.015 | 0.036 | -0.424 | 0.672 |
| Reptiles | -0.017 | 0.042 | -0.392 | 0.696 |
| Birds | 0.003 | 0.006 | 0.342 | 0.733 |
| Mammals | -0.054 | 0.049 | -1.113 | 0.268 |
| Vertebrates | 0.019 | 0.024 | -0.795 | 0.428 |
| Plants | 0.007 | 0.006 | 1.162 | 0.247 |
